# Using phylogenetic relationships to assess conservation priorities for birds in India

**DOI:** 10.1101/2025.08.04.668436

**Authors:** Archita Sharma, Naman Goyal, Dhyey Shah, V. V. Robin

**Affiliations:** Department of Biology, Indian Institute of Science Education and Research Tirupati, Tirupati, Andhra Pradesh, India; Department of Zoology, The Maharaja Sayajirao University of Baroda, Vadodara, Gujarat

**Keywords:** EDGE score, evolutionary distinctiveness, India, Ornithology

## Abstract

In the midst of the ongoing biodiversity crisis, setting conservation priorities has moved beyond the conventional approaches. The need to incorporate phylogeny-based metrics that capture evolutionary rarity with extinction risk for prioritization of global conservation efforts has been widely recognized. Such metrics are even more important in megadiverse countries of the Global South, where conservation spending is limited despite high conservation value. Here, we reassess global avian threatened evolutionary history with an updated phylogeny, using the EDGE metric. The EDGE metric or score combines evolutionary distinctiveness of a species with its extinction risk. We focus on India to identify species and regions that represent high avian EDGE scores. We find that the Bengal Florican (*Houbaropsis bengalensis*) had the highest median EDGE score of 18.83 MY (millions of years). To identify spatial conservation priorities, we mapped avian EDGE scores at a 10 km x 10 km scale. Northeastern India is exceptional in its richness of evolutionary unique and threatened species. We also scored IBAs and PAs, finding that IBA, as well as PA, with the maximum EDGE score, was Intaki or Ntangki National Park in Nagaland, accumulating ∼250 MY of threatened evolutionary history in India. Habitats of the top 5% of EDGE species in India are currently not adequately protected. Our study offers a preliminary basis for a quantitatively informed and phylogeny-driven conservation prioritization approach for researchers, practitioners, and policymakers to safeguard threatened evolutionary history of birds in India.

## 1. Introduction

Increasing evidence suggests that many species face a heightened risk of extinction across the globe (Pimm et al. 2014; Ceballos and Ehrlich 2023), with the growing need to identify species and areas that need to be prioritized for conservation action and resource allocation. Several studies have argued that the biodiversity value of an area is better represented by the amount of evolutionary or phylogenetic diversity it hosts rather than just its species richness (Vane-Wright, Humphries, and Williams 1991; Faith 1992; Crozier 1997; Pio et al. 2011). Species vary in the evolutionary history that they represent across the tree of life. Moreover, species with fewer close relatives are more threatened when compared to species with many close relatives (Purvis et al. 2000). The usage of the EDGE metric (Isaac et al. 2007) representing threatened evolutionary history of a species by combining its evolutionary distinctiveness (ED) with its extinction risk or global endangerment (GE) has gained momentum in recent years. It has been calculated for several taxonomic groups - amphibians (Isaac et al. 2012), birds (Jetz et al. 2014), corals (Huang 2012), mammals (Isaac et al. 2007), and all tetrapods (Gumbs et al. 2018). More recently, an updated EDGE2 metric has been developed, which is more robust to uncertainty in extinction risk data and phylogenetic data (Gumbs et al. 2023).

EDGE2 is a product of ED2, capturing the irreplaceability of a species, and GE2, capturing the vulnerability of a species. One of the key advances in the EDGE2 metric from the EDGE metric is incorporating phylogenetic diversity (PD) complementarity in the ED2 calculation (Gumbs et al. 2023). Conceptually, ED2 captures the influence of the extinction of close relatives in the future on the focal species’ irreplaceability (Steel, Mimoto, and Mooers 2007). EDGE2 metric or scores (EDGE2 scores hereafter) represent threatened evolutionary history, with scores representing evolutionary history in millions of years (MY) to be lost if not conserved (Gumbs et al. 2023). EDGE2 scores have been calculated globally for birds (McClure et al. 2023) and for broader taxonomic groups such as tetrapods (Pipins et al. 2024) and jawed vertebrates (Gumbs et al. 2024). All of these studies sourced the data for birds from the same global avian phylogenetic tree (Jetz et al. 2012, 2014), which followed the taxonomy of BirdLife v3 world list (June 2010) and IOC World Bird list v2.7 (Dec 29, 2010) (Gill, Wright, and Donsker 2010), resulting in 9993 recognized species. This phylogenetic tree has been widely used as a backbone tree for many large-scale ornithological studies (Pigot et al. 2020; Delhey et al. 2023; Neate-Clegg et al. 2023); however, it incorporates genetic data for only 66% of the total species (6663 out of 9993) and relies on phylogenetic imputation or taxonomic constraints for the missing species. Phylogenetic imputation does not accurately capture the true ED values or rank of missing species (Weedop et al. 2019). A decade later, a more recent global bird tree is now available, which incorporates information from 166 bird phylogeny studies post-2012, adding genetic data for 9189 out of 11017 species (83%) (McTavish et al. 2025). There are 2377 branches in this phylogenetic tree that capture different phylogenetic relationships for various species compared to the previously published global phylogeny (McTavish et al. 2025). Re-assessing the global avian EDGE2 scores using an updated phylogeny allows us to compare and contrast values from previous efforts and further investigate regional-scale patterns more accurately.

India is among the top 10 countries (ranked 3) projected to suffer the highest evolutionary loss (measured as a loss of Phylogenetic Diversity) due to land-use-driven species extinctions (Chaudhary, Pourfaraj, and Mooers 2018). Minimizing future loss of India’s threatened evolutionary history should be a top priority. India is a large tropical country with four global biodiversity hotspots, a network of protected areas (PAs) and Important Bird Areas (IBAs), and distinct biogeographic zones. Its rich biodiversity can be traced back to vicariance events like the Gondwana break-up, as well as into-India dispersal events post-Gondwana break-up (Karanth 2021), resulting in many evolutionary distinct lineages. While the idea of setting conservation priorities using evolutionary or phylogenetic information has been around for decades, it had not really been picked up in India until a handful of very recent studies (Bharti et al. 2021; Shooner et al. 2018; Gopal et al. 2023; Goyal et al. 2025; Manish and Pandit 2018).

For birds in India, to set conservation priorities, country-scale studies have relied on species distribution data and citizen science approaches. Evaluation of the status of birds in India using detection and non-detection data from eBird identified conservation priorities for 942 bird species as part of the comprehensive State of India’s Birds (SOIB) 2023 report (Viswanathan et al. 2025). Another study calculated the Area of Habitat (AOH) for 99 of India’s small-range threatened bird species and compared PA coverage (Warudkar et al. 2022). AOH represents habitat available to a species within its range, and can complement range maps by calculating potential occupancy and removing false presences (Brooks et al. 2019). For birds, AOH maps are available at a global scale, improving the accuracy of species distributions and conservation planning (Lumbierres et al. 2022). Assessing the PA coverage using AOH of the top EDGE species in India would allow us to understand how well protected India’s most evolutionary unique and threatened species are.

In this study, using the EDGE2 (hereafter EDGE) framework, we proposed to a) reassess global avian EDGE scores and b) map avian EDGE scores in India at a 10 km x 10 km scale, identifying species and areas (broad biogeographic regions, IBAs, and PAs) with high scores. For India’s top EDGE species, we plan to use the already available fine-scale Area of Habitat (AOH) maps to calculate the proportion of area inside and outside the Protected Areas (PAs) in India. We expect the ED scores and EDGE scores, and rankings to change across the global avian tree of life with the use of an updated phylogeny. Global EDGE studies at broader spatial scales (96.5 km x 96.5 km) have identified the Himalayas and southern Western Ghats to be important regions in India with respect to EDGE species richness for tetrapods (Pipins et al. 2024). Both of these are global biodiversity hotspots. At finer spatial scales (10 km x 10 km), we expect other underrepresented areas in the North Western Ghats, Satpuras, and Eastern Ghats to also show patterns. We also expect a significant proportion of the AOH area to be outside PAs for India’s top EDGE bird species.

## 2. Methods

### 2.1 EDGE Scores for birds - Global

We applied the EDGE protocol (Gumbs et al. 2023) to re-evaluate global EDGE prioritization for birds worldwide. We used a recently published phylogeny for global aves comprising 11017 extant and extinct species, following eBird/Clements 2023 taxonomy (McTavish et al. 2025). For this study, we follow the taxonomy used in eBird/Clements Checklist v2023 (Clements et al. 2023). We matched this taxonomy with HBW and BirdLife International Checklist v9 (2024) to get IUCN Red List data. We reconciled the two lists and assigned the red list categories to synonyms (the same species is listed under different names in the two lists). For species that were split/lumped in Clements Checklist v2023 but not in HBW Checklist v9, and were not exact matches between the two lists, were kept as NE (Not Evaluated). The recently compiled AviList (Rheindt et al. 2025) also includes many species as NE owing to taxonomic mismatch with BirdLife International (used by IUCN for the Red List).

We calculated the GE2 scores or the probability of extinction, *pext*, using the GE2 simulation code (https://github.com/rgumbs/EDGE2) in R version 4.4.1. In this approach, the *pext* value is assumed to vary continuously between species and not jump as we move between discrete IUCN threat categories (Gumbs et al. 2023). The *pext* varies between 0 and 1, where 0 is completely safe, and 1 is extinct. Each of the five IUCN Red List categories is assigned a predefined median value within this range following Gumbs et al. 2023 (CR = 0.97, EN = 0.485, VU = 0.2425, NT = 0.12125, LC = 0.0606025). A smooth quartic curve is fit based on realistic constraints with *pext* bounded between 0.0001 (safe) and 0.9999 (almost certain to go extinct), passing through the five median values corresponding to each IUCN Red List category (Gumbs et al. 2023). The portion of the distribution is equally assigned to each category. For each species, *pext* or GE2 scores are then drawn corresponding to the IUCN Red List category from the curve. We also included extinct species by setting the value of *pext* to 1, thereby removing their influence on EDGE scores (Gumbs et al. 2023). Further, we included the Data Deficient/Not Evaluated (DD/NE) category by assigning the value of *pext* to the median of the entire range (0.0001-0.9999, median = 0.232) following Gumbs et al. (2023). The median *pext* values used for each IUCN category and the count of bird species for each category are mentioned in Appendix A, Table A1.

To capture phylogenetic uncertainty, it is suggested to calculate *pext* values a large number of times for each species (100 < n ≤ 1000 trees) (Gumbs et al. 2023). We used a random sample of 100 phylogenetic trees using a randomly sampled node age for each dated node calibration available at McTavish Lab’s GitHub repository (McTavish 2025). We repeated our EDGE calculations across 100 phylogenetic trees. For each tree, a new *pext* value was selected from the bounds of uncertainty for the given IUCN Red List category (Figure A1). For 11017 species, we calculated the median, 2.5th, and 97.5th percentiles of ED and EDGE scores, along with TBL (Terminal Branch Length) and *pext* values. See the GitHub repository (https://github.com/rgumbs/EDGE2) for functions to calculate EDGE scores. The final EDGE score of a species is taken as the median of EDGE scores across 100 trees, to account for uncertainty in phylogenetic position and probability of extinction. We calculated ED and EDGE for each avian order (n=41) and compared our results to a previous study (McClure et al. 2023). The orders might not be entirely comparable owing to the different versions of taxonomy used in the two studies. Between the two studies, only 36 orders were common, while 5 were unique to our study as a result of updates in avian taxonomy - Rheiformes, Apterygiformes, Casuariiformes, Galbuliformes, and Tinamiformes.

### 2.2. EDGE Scores for birds - India

We used the most updated species list of India, which contained 1373 species (Praveen and Jayapal 2025). We reconciled this list with the Clements 2023 taxonomy to get a final list of 1372 species. We created a subset of the global EDGE scores of 1372 species found in India. To identify the most evolutionarily distinct and threatened species in India, we ranked all 1372 species based on their median EDGE scores in descending order. We identified the top 5% of species by calculating the 95th percentile of the median EDGE score distribution. Species with EDGE scores greater than or equal to this threshold were classified as the top 5% EDGE species in India (n=69) (Figure A4).

To get the distribution ranges of birds found in India, we downloaded digitized maps compiled by Birdlife International for global birds (following HBW / BirdLife Taxonomic Checklist v9 taxonomy, Version 2024.2). We fixed geometries for invalid layers and merged the valid layers to clip this valid dataset with the India shapefile. We then made a crosswalk between Clements 2023 taxonomy (Clements et al. 2023) and BirdLife taxonomy (HBW and BirdLife International 2024) to get shapefiles for 1248 species of birds found in India as per Clements 2023 taxonomy. Spatial analysis is limited to 1248 species, excluding Narcondam Hornbill (not spatially aligned with Narcondam island) (*Rhyticeros narcondami*), Buff-chested Babbler (*Cyanoderma ambiguum*), and Malabar Starling (*Sturnia blythii*). We then transformed the spatial distribution of species in India into a presence-absence matrix at the scale of 10 km x 10 km (∼0.1 degree) equal area grids using the letsR package (Vilela and Villalobos 2015). The BirdLife range maps may not be accurate at such a fine spatial scale, but we decided on this scale because the protected area size in India varies anywhere between 1 square kilometer for some Islands to a few larger than 5000 square kilometers (Ghosh-Harihar et al. 2019). We then calculated the total EDGE score for each grid by summing the EDGE score of each species found in that particular grid. The resulting matrix was spatially mapped in R version 4.4.1 to visualize the spread of EDGE scores across India. Linear regression analysis examining the spatial relationship between the EDGE score and species richness was also done (Figure A6).

We also downloaded the IBA shapefiles of India (BirdLife International 2024) and PA shapefiles of India (https://corridorcoalition.org/), which represent a large number of the known PAs of India, while the boundaries of other PAs are either undocumented or not available in the public domain. We then scored each IBA and PA by extracting the scores of all 10 km x 10 km grids falling in each IBA and PA. For each IBA and PA, the final EDGE score was decided by the grid with the maximum EDGE score within its area, respectively.

India has ten zones: Coasts, Deccan Plateau, Deserts, Gangetic Plains, Himalayas, Islands, Northeast India, Semi-arid, Trans-Himalayas, and Western Ghats (Rodgers and Panwar 1988). We downloaded the shapefiles for biogeographic zones in India (Vattakaven et al. 2016) and identified the top five IBAs and PAs with the highest EDGE scores from each biogeographic zone. Each IBA and PA was assigned to a single biogeographic zone based on the zone with which they had the greatest spatial overlap in area.

For the top 5% EDGE bird species in India, we downloaded already published AOH maps at a resolution of 100 meters (Lumbierres et al. 2022). We calculated the AOH area inside and outside the PAs in India. We used zonal statistics in QGIS v3.22.4 to get pixel-scale statistics from species-specific raster layers and the PAs vector layer.

## 3. Results

In this study, we re-evaluated EDGE scores for all birds across the globe using the McTavish et al. (2025) phylogeny, which follows a relatively newer taxonomy (ebird/Clements 2023 taxonomy), and compared the scores to those from a previous study (McClure et al. 2023), which used Jetz et al. (2012) phylogeny. We also created a subset of the species in India to understand fine-scale spatial patterns in the distribution of EDGE scores across the country and identify species and areas with high EDGE scores.

### 3.1. EDGE scores for birds - Global

Most avian orders show similar EDGE scores between this study and McClure et al. (2023) (Figure 1). The most notable difference is in Order Struthioiformes, which has an EDGE score of 7.1 MY in this study compared to an EDGE score of 0.49 MY in McClure et al. (2023), suggesting increased conservation priority. The Orders with the highest median EDGE scores were Eurypygiformes (includes Kagu [*Rhynochetos jubatus*] and sunbittern [*Eurypyga helias*]), Struthioniformes (includes Ostriches), and Opisthocomiformes (includes Hoatzin [*Opisthocomus hoazin*]) (Figure A2). ED scores show more variability between the two studies owing to the use of different phylogenetic trees (Figure 1). The orders with the greatest median ED scores were Opisthocomiformes, Leptosomiformes, and Struthioniformes (Figure A3).

**Figure 1.**
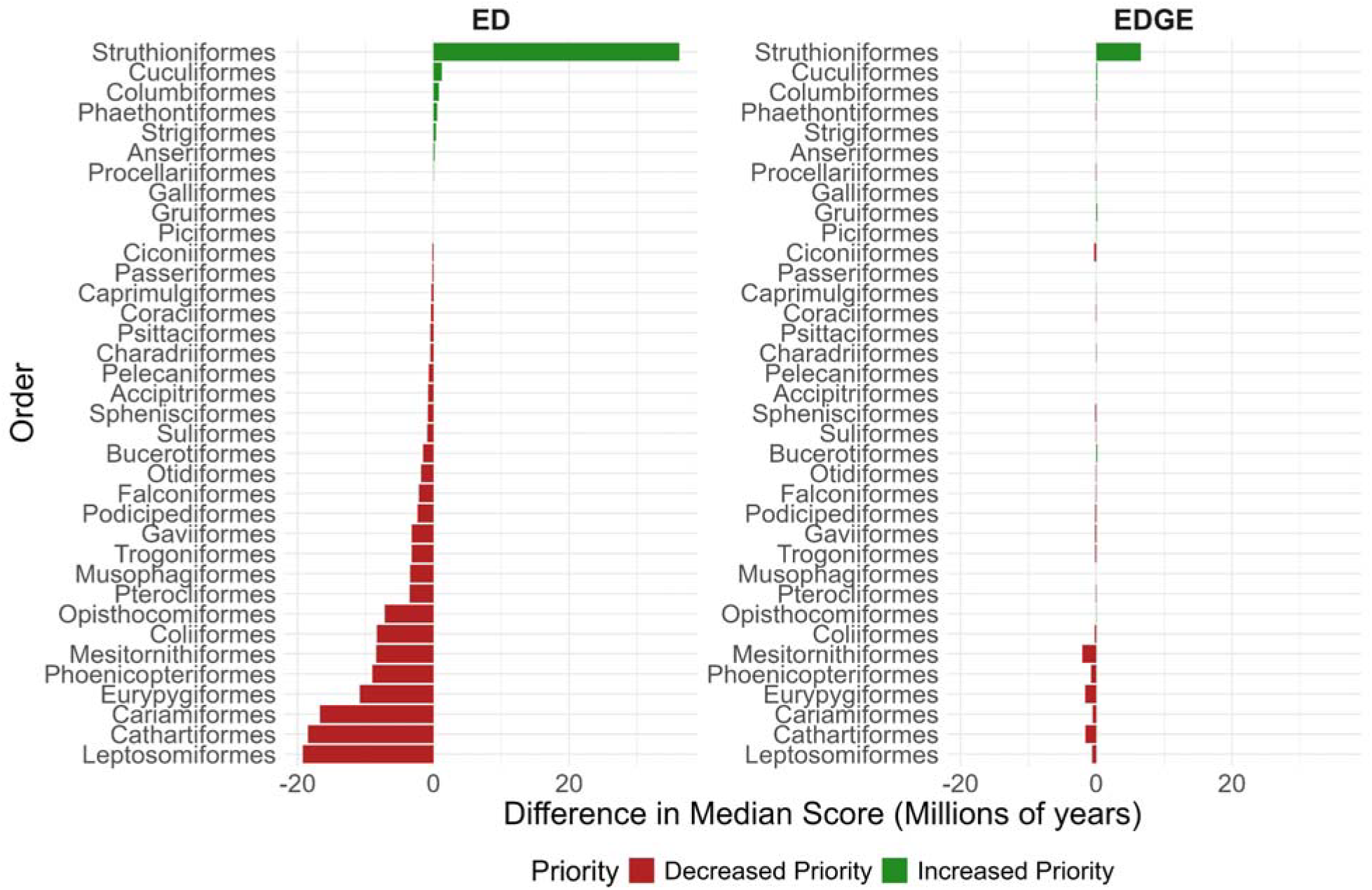
Change in median ED score and EDGE score for 36 common avian orders between this study and McClure et al. (2023). Increased Priority implies a higher ED or EDGE score in this study when compared to McClure et al. (2023).

In this study, among the extant species of birds, Maleo (*Macrocephalon maleo*) had the highest median EDGE score (26.159 MY), and Japanese Pygmy Woodpecker (*Yungipicus kizuki*) had the lowest EDGE score (0.000037 MY). For the global top 30 extant species with the highest median EDGE scores, we also document changes in conservation priority based on the results of this study when compared to McClure et al. (2023) (Figure 2). Increased priority implies that the species is ranked higher (higher EDGE score) in this study. For instance, the Somali Ostrich (*Struthio molybdophanes*) had a score of 2.11 MY and a global rank of 402 in (McClure et al. 2023), but in our study, it had a score of 11.1 MY and a global rank of 22. Species like Somali Ostrich (*Struthio molybdophanes*), Himalayan Quail (*Ophrysia superciliosa*), Polynesian Storm-Petrel (*Nesofregetta fuliginosa*), and Jamaican Pauraque (*Siphonorhis americana*) jumped up by more than 100 ranks, suggesting increased conservation priority than previously estimated. Species like the Rufous Scrub-bird (*Siphonorhis americana*), Giant Ibis (*Pseudibis gigantea*), Dwarf Tinamou (*Taoniscus nanus*), and Noisy Scrub-bird (*Atrichornis clamosus*) had slightly lower ranks in our analysis than previously anticipated, suggesting decreased conservation priority; nevertheless, these remain high-scoring EDGE species. Several extant species (n=339) were also not common between the two lists, and ranks could not be compared for those.

**Figure 2.**
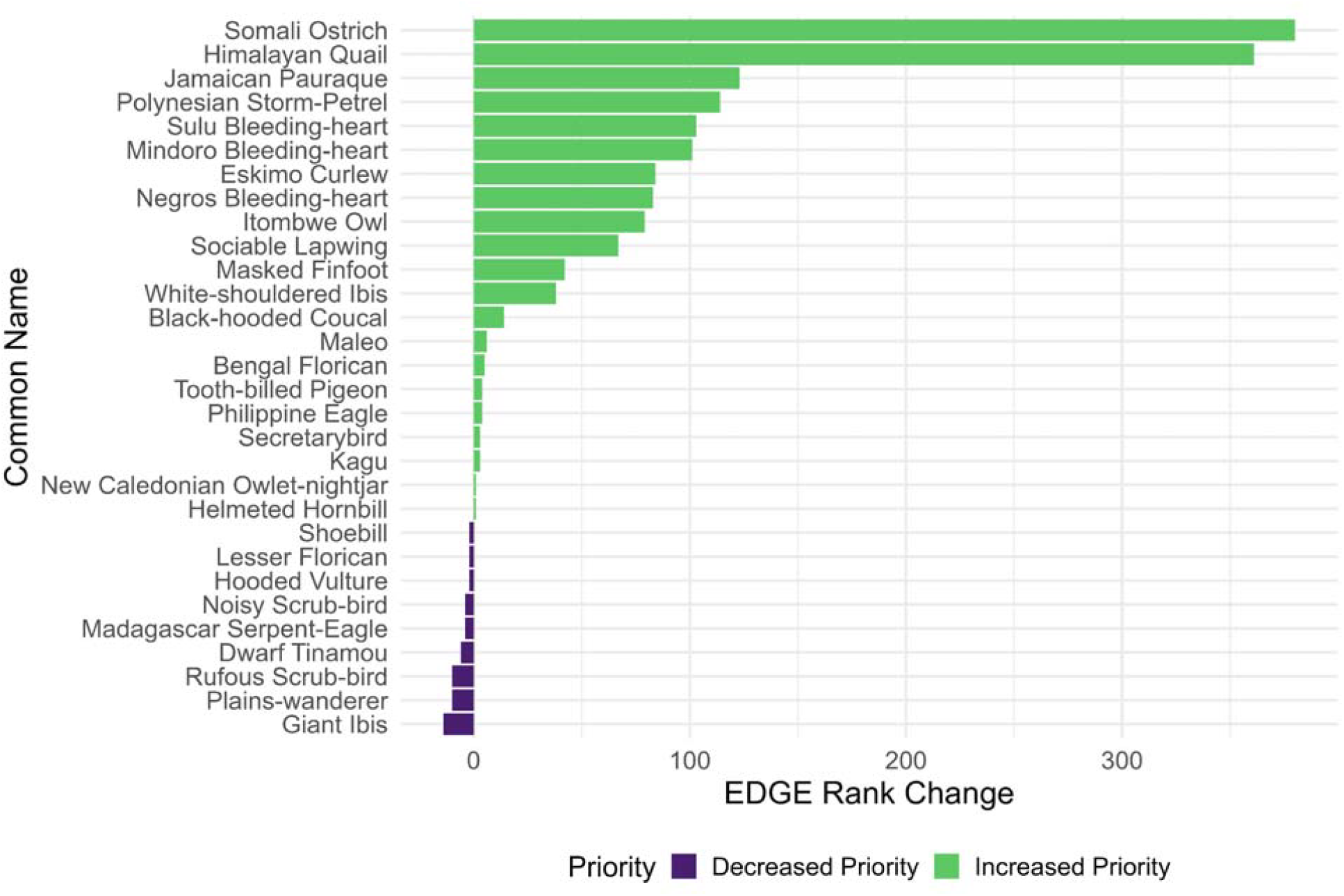
EDGE rank changes between this study and McClure et al. (2023) for the top 30 extant EDGE species in this study, which are common with the McClure et al. (2023) species list. Increased Priority implies that the species has a higher rank (higher EDGE score) in our study.

A notable number of bird species (310 out of 11017) remain Data Deficient or Not Assessed (DD/NE). Accurate global assessments of DD/NE species, mostly arising from splits, lumps, and shuffles in avian taxonomy, could help in arriving at more precise estimates of extinction risks and EDGE scores for these species.

### 3.2 EDGE scores for birds - India

As per the Indian Birds Checklist v9, 1372 species are found in India (including historical and vagrant species, as per Clements 2023 taxonomy). Summing EDGE scores for all 1372 species, overall, India represents 639 MY of threatened evolutionary history. Among the species found in India, Bengal Florican (*Houbaropsis bengalensis*) had the highest median EDGE score of 18.83 MY, and Brown-capped Pygmy Woodpecker (*Yungipicus nanus*) had the lowest median EDGE score of 0.0000679 MY. Hodgson’s Frogmouth (*Batrachostomus hodgsoni*) had the highest median ED score of 45.21 MY.

#### 3.2.1 Top 5% EDGE species

The top 5% evolutionary distinct and globally endangered species found in India (n=69) included mostly critically endangered and endangered species on the IUCN list (Figure 3). However, 7 of India’s top 5% EDGE species are classified as least concern: Hodgson’s Frogmouth (*Batrachostomus hodgsoni*), Osprey (*Pandion haliaetus*), Sri Lanka Frogmouth (*Batrachostomus moniliger*), Red-throated Diver (*Gavia stellata*), Great Eared-Nightjar (*Lyncornis macrotis*), Jack Snipe (*Lymnocryptes minimus*), and Crab-Plover (*Dromas ardeola*). While 31 of India’s top 5% EDGE species have been classified as ‘high’ conservation priority by the State of India’s Birds 2023 framework (Viswanathan et al. 2025), there are 4 species that are classified as ‘low’ conservation priority.

**Figure 3.**
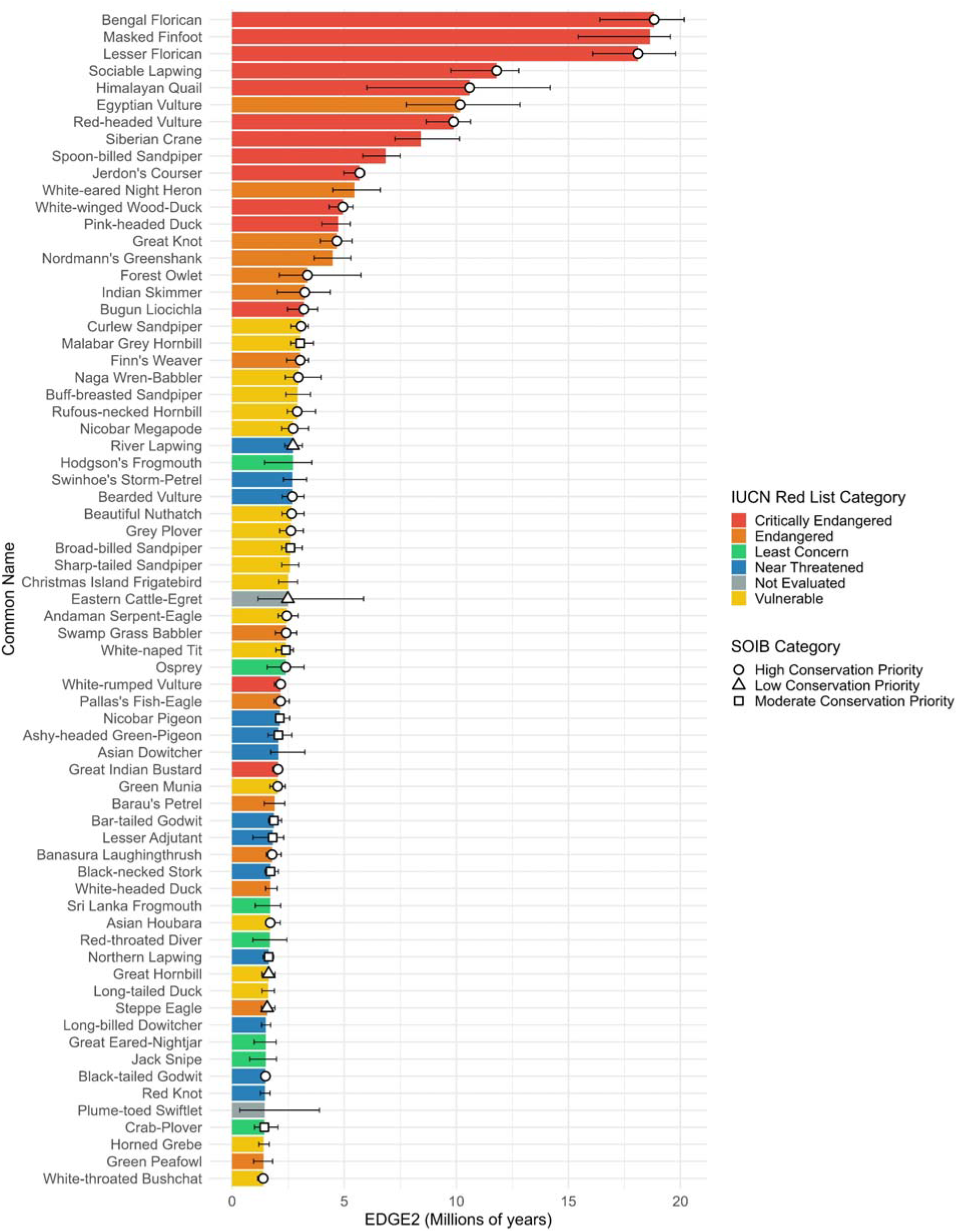
Top 5% species of birds (n=69) in India with the highest median EDGE scores (including historical and vagrant records).

This list also includes species with only historic records (n = 5) from India: Masked Finfoot (*Heliopais personatus*), Himalayan Quail (*Ophrysia superciliosa*), Pink-headed Duck (*Rhodonessa caryophyllacea*), Barau’s Petrel (*Pterodroma baraui*), and Green Peafowl (*Pavo muticus*). Similarly, species with vagrant records (n = 11) from India are also included in the top 5%: Siberian Crane (*Leucogeranus leucogeranus*), Spoon-billed Sandpiper (*Calidris pygmaea*), White-eared Night Heron (*Oroanassa magnifica*), Nordmann’s Greenshank (*Tringa guttifer*), and others. A total of 11 species are endemic to India - Himalayan Quail (*Ophrysia superciliosa*), Jerdon’s Courser (*Rhinoptilus bitorquatus*), Forest Owlet (*Athene blewitti*), Bugun Liocichla (*Liocichla bugunorum*), Malabar Grey Hornbill (*Ocyceros griseus*), Naga Wren-Babbler (*Spelaeornis chocolatinus*), Nicobar Megapode (*Megapodius nicobariensis*), Andaman Serpent-Eagle (*Spilornis elgini*), White-naped Tit (*Machlolophus nuchalis*), Green Munia (*Amandava formosa*) and Banasura Laughingthrush (*Montecincla jerdoni*). Out of 69 species, 55.4% (n=36) are migratory and 44.6% (n=33) are non-migratory or resident species. We also identified the top 5% EDGE species after excluding historical and vagrant records (Figure A5).

#### 3.2.3 Spatial Analysis

To identify spatial conservation priorities, we mapped EDGE scores of 1248 bird species at a 10 km x 10 km scale across India. Geographically, the highest EDGE scores were concentrated in northeastern India, the Himalayas, and the Western Ghats (Figure 4). Parts of Central India (Satpuras) and northern Eastern Ghats also show high EDGE scores. Linear regression analysis examining the spatial relationship between the EDGE score and species richness revealed that they are positively correlated (R^2^=0.745, Figure A6). Calculated residual values across India’s biogeographic zones revealed that the Semi-Arid, Gangetic Plain, Western Ghats, and Deccan Peninsula zones had higher EDGE scores than the model prediction. These zones have higher EDGE scores than expected given their species richness, revealing that they host higher than expected threatened evolutionary history for birds in India.

**Figure 4.**
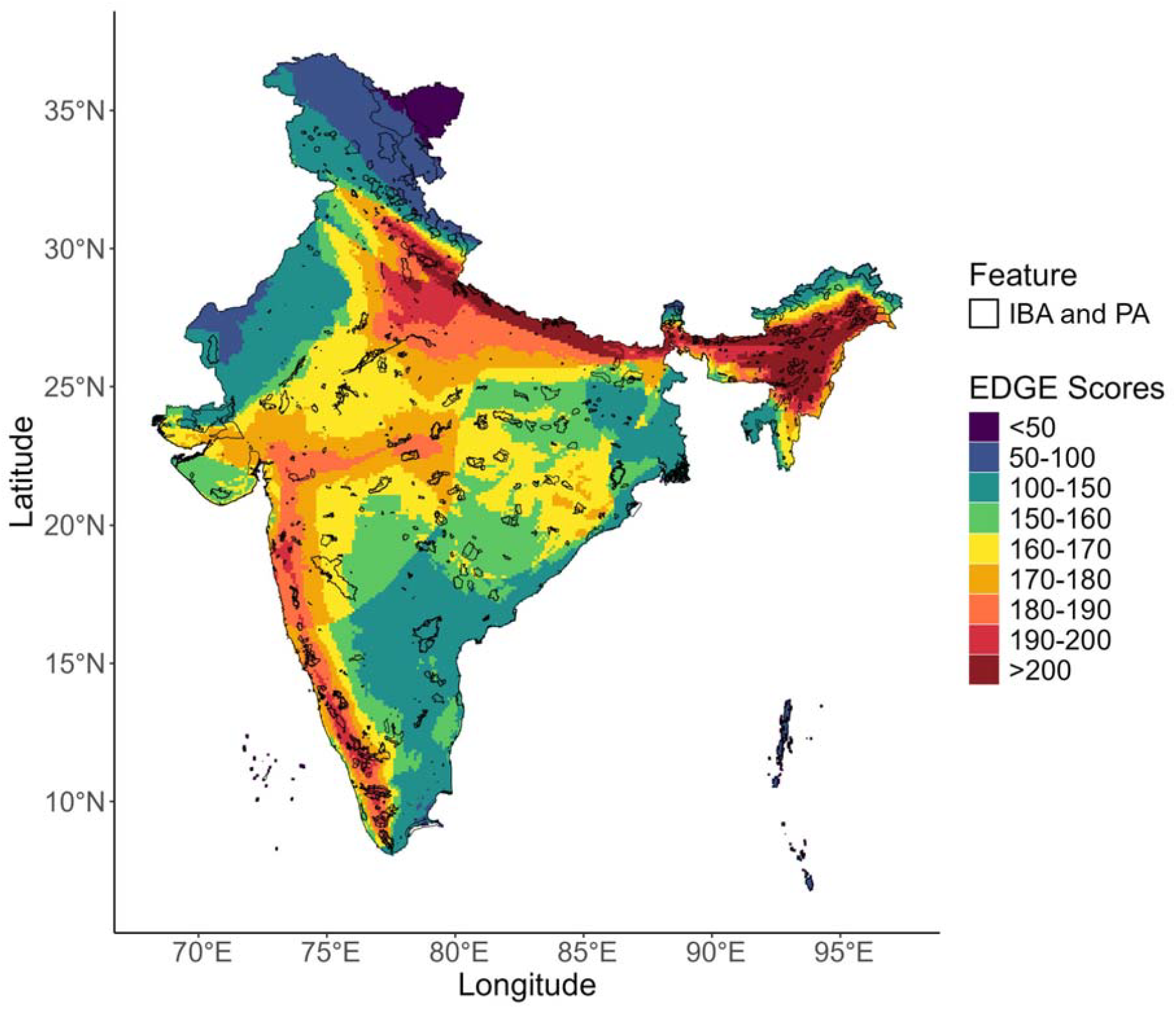
Spatial distribution of EDGE scores (10 km x 10 km grids) along with IBA and PA boundaries

We also calculated EDGE scores for 551 IBAs and 501 PAs in India. In India, IBA, as well as PA with the maximum EDGE score, was Intaki or Ntangki National Park in Nagaland. Additionally, we identified the five top-scoring IBAs (Figure 5) and PAs (Figure A7) for each of India’s biogeographic zones. The highest-scoring IBAs were Thane Creek in the Coasts zone, Bhimashankar Wildlife Sanctuary in the Deccan Peninsula Zone, Banni Grassland and Chhari Dhand in the Desert Zone, Pilibhit Tiger Reserve in the Gangetic Plain Zone, Eaglenest Wildlife Sanctuary in the Himalaya Zone, Mount Harriet National Park (3 more with same score) in the Islands Zone, Intaki National Park in the Northeast Zone, Sur Sarovar Bird Sanctuary in the Semi-Arid Zone, Kangchendzonga National Park and Biosphere Reserve in the Trans-Himalayan Zone, and Kottiyoor Reserve Forest (2 more with the same score) in the Western Ghats.

**Figure 5.**
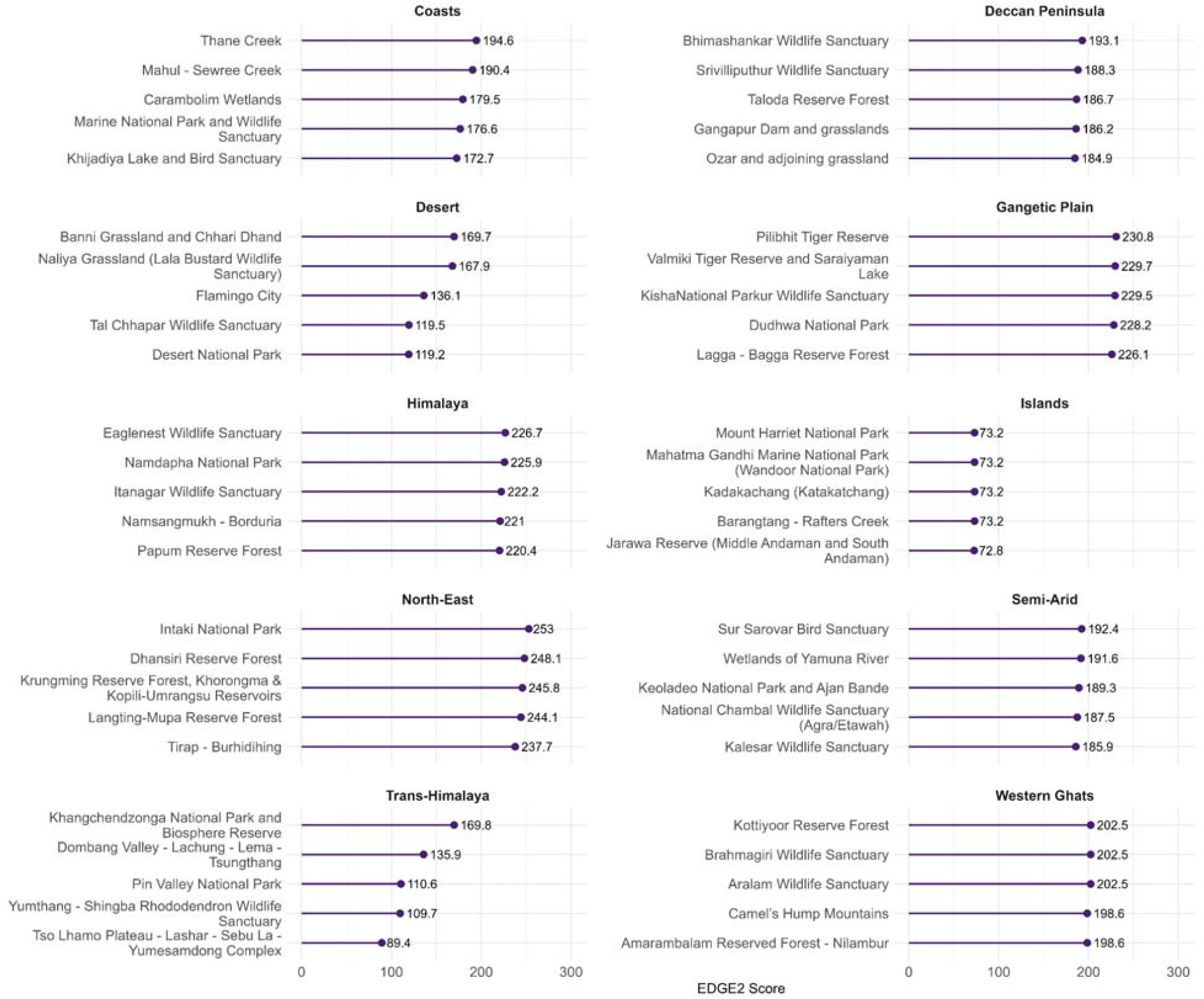
Top IBAs (Important Bird Areas) with the highest EDGE scores in each of India’s 10 biogeographic zones.

Out of 69 species, we were able to access AOH for 64 species, excluding Sri Lanka Frogmouth (*Batrachostomus moniliger*), Swinhoe’s Storm-Petrel (*Hydrobates monorhis*), Plume-toed Swiftlet (*Collocalia affinis*), and Crab-Plover (*Dromas ardeola*), since these were not available. Out of 64 species, some species did not have AOH within the bounds of India, possibly because many of these species have vagrant records from India. For 53 out of 69 species (in the top 5%), we did a more fine-scale spatial analysis using globally calculated AOH maps and Protected Area shapefiles. Our calculations of AOH area inside and outside PAs revealed that only 12 out of 53 species had more than 10% of their AOH area protected (Figure 6). This includes Forest Owlet, with almost 50% of the AOH area of the is protected despite being range-restricted, which is the highest among the top 5% EDGE species. However, most of India’s top 5% EDGE species have only a small proportion of their AOH range currently protected.

**Figure 6.**
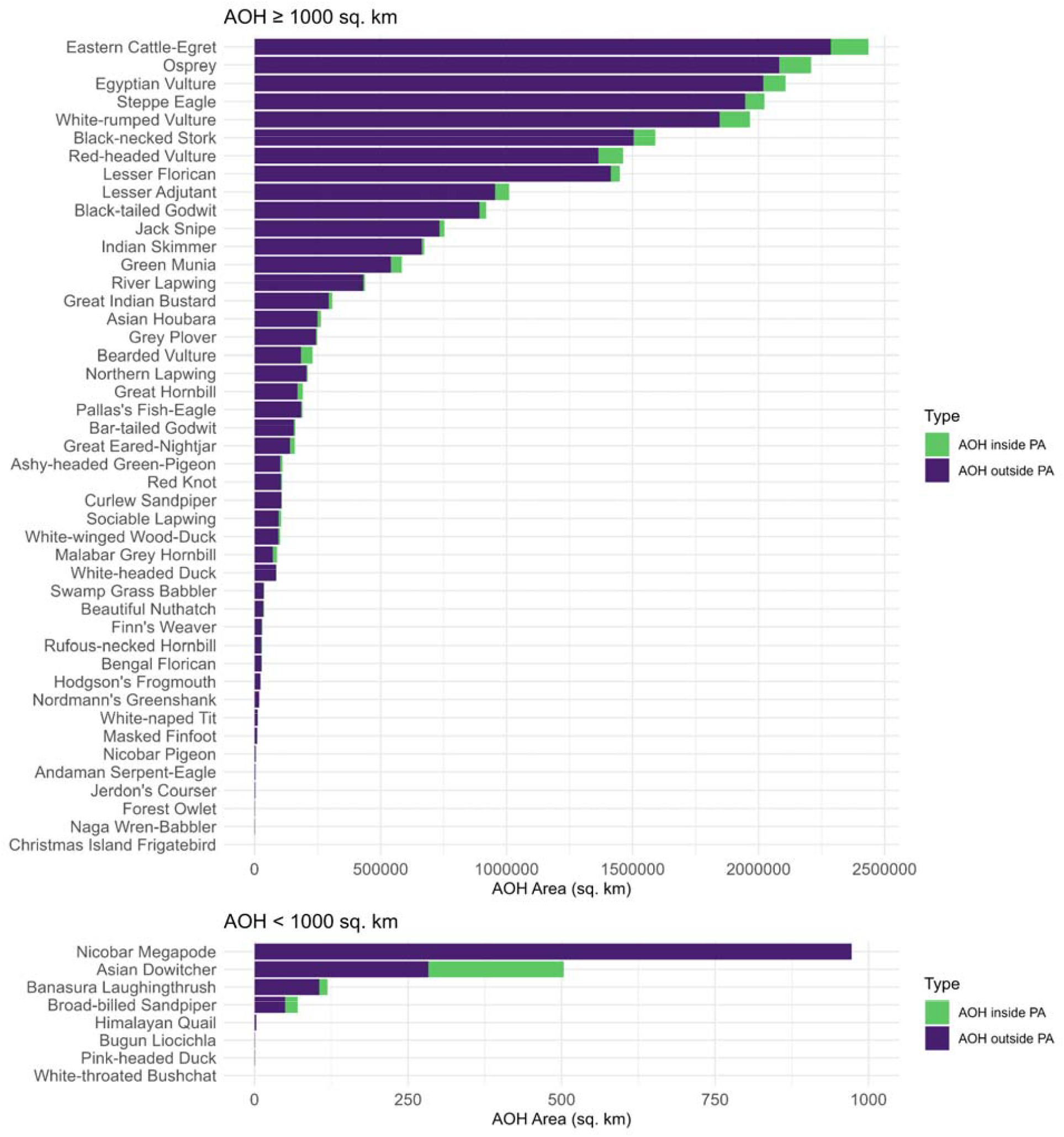
AOH (Area of Habitat) in square kilometers for the top 5% EDGE species and proportion of area outside and inside PA (Protected Areas). It includes 53 of 69 species for which AOH maps could be obtained and were spatially within India’s geographic boundaries.

## 4. Discussion

Global phylogenies that include all members of a clade are an important resource for evolutionary inference and setting phylogenetically informed conservation priorities. We re-evaluated the threatened evolutionary history of global birds, and we identified several species that require increased conservation attention. Additionally, at a smaller scale – within India, we identified locations that host disproportionate numbers of such species.

Our analysis differed from previous similar exercises (Pipins et al. 2024; McClure et al. 2023; Gumbs et al. 2024) in using a different and updated phylogeny (McTavish et al. 2025). At the scale of avian orders, Struthioniformes show a considerable difference in median EDGE scores between the two studies, with the Somali Ostrich moving up in the EDGE species rankings and needing increased conservation priority. This difference is due to a change in the taxonomy and phylogenetic position of Struthioniformes between the Jetz et al. (2012) and McTavish et al. (2025) phylogenetic trees. Significantly, the McTavish phylogeny (McTavish et al. 2025) is created on a platform that permits new genetic information to be regularly updated. This provides us with an opportunity to update the conservation significance of species as new genetic data becomes available.

Our study finds that the species with the highest EDGE score in India is the Bengal Florican (*Houbaropsis bengalensis*). Bengal Floricans have their breeding ranges in grasslands, while their non-breeding ranges are often in agricultural-floodplain landscapes outside PAs (Jha et al. 2018). Conservation of grassland habitats and floodplains is crucial for the conservation of Bengal Floricans (Thakur et al. 2024; Packman et al. 2014). Their appropriate habitat (i.e., AOH) is only 2.18% of their published range (Warudkar et al. 2022). Additionally, only ∼7% of the species’ AOH area is currently protected. This clearly highlights the conservation challenge of a grassland specialist that is evolutionarily distinct, has very little habitat in its range, and has very little of its habitat protected. We also identified India’s top 5% EDGE species, many of which are either critically endangered, endangered, or vulnerable. Additionally, many of them are classified with ‘high’ conservation priority status in the SOIB 2023 report based on range sizes, abundance, and long-term change assessments (Viswanathan et al. 2025). 24 of the top EDGE species are also included in either Appendix 1 or Appendix 2 of the CITES Appendix. Our study reinforces the need to prioritize conservation of these threatened species, which are also evolutionary unique in India.

However, seven of the top EDGE species are of the Least Concern category as per the IUCN Red List, and four species are classified with low conservation priority status as per the SOIB 2023 report (Viswanathan et al. 2025). These species may not be as threatened or declining in abundance trends currently; however, they represent high ED scores, which means that the expected contribution of these species to unique phylogenetic diversity in the future would be high because they have very few close relatives, and these close relatives currently face high extinction probabilities. To simplify, in a future scenario where all its close relatives were to perish, this species would represent a single most unique evolutionary lineage, leading to high ‘expected distinctiveness’ (Gumbs et al. 2023). Therefore, we highlight a set of species in India that are important for conservation prioritization from a phylogenetic perspective. It is also worth noting that species like Eastern Cattle Egret (*Bubulcus coromandus*) have made it into the top 5% due to a highly variable *pext* since the IUCN status remains unassessed post its split from Western Cattle Egret (*Bubulcus ibis*) into a newly recognized species.

In India, the Himalayas and Western Ghats have high species richness and endemism, and it was expected that these areas would harbour grids with high EDGE scores. Grids with high EDGE score were also scattered through the Gangetic plains, northern Western Ghats, Satpuras, and the northern Eastern Ghats in peninsular India. In peninsular India, these mountain ranges are important biogeographic units showing species and subspecies turnovers for birds across topographic and climatic barriers (Ramachandran et al. 2017). The spatial distribution of EDGE scores in the peninsular part of India is similar to patterns of phylogenetic diversity of birds observed in a recent study (Goyal et al. 2025). It is not surprising that northeastern India, part of the Indo-Burma biodiversity hotspot – one of the most biodiverse regions in the world (Price 2012) has emerged to be one of the most important areas in India with the most evolutionarily unique and threatened species. The IBA and PA in India with the highest EDGE score was in this region – the Intaki or Ntangki National Park (INP) in Nagaland. A recent study estimated an 11% loss of forest cover in INP between 1999-2017 based on remote sensing data (Liezietsu et al. 2022), apart from poaching pressures in the broader region (Velho, Karanth, and Laurance 2012), indicating further significant threats in the region.

However, it is also important to note that the spatial spread of EDGE scores was highly correlated with the spatial spread of avian species richness in India. Biogeographic zones in India that host higher than expected threatened evolutionary history for birds, given the species richness, were - Semi-Arid, Gangetic Plain, Western Ghats, and Deccan Peninsula. This highlights the importance of these regions in encapsulating high amounts of threatened evolutionary history in India, by accumulating species that are evolutionarily unique, despite having overall lower amounts of avian species richness when compared to other biogeographic regions.

We also assessed the proportion of habitat within protected areas for the top 5% EDGE species in India. Our study found that most species with high EDGE scores had less than 10% of their Area of Habitat (AOH) protected. For species whose entire geographic area is <1000 square kilometers, it is recommended to have 100% complete coverage by protected areas (Venter et al. 2014). Five of the top 5% EDGE species in our analysis - Nicobar Megapode (*Megapodius nicobariensis*), Banasura Laughingthrush (*Montecincla jerdoni*), Himalayan Quail (*Ophrysia superciliosa*), Bugun Liocichla (*Liocichla bugunorum*), and Pink-headed Duck (*Rhodonessa caryophyllacea*) are endemic to India and have an AOH area <1000 square kilometers, and additionally have at most 25% of this area covered by PAs. Our study highlights the scope to expand current protected area coverage in India to encompass the spatial habitats and ranges of such evolutionary unique and threatened bird species.

We note that our study is limited by the resolution of available species distribution data (from BirdLife International) at a low resolution, while our study is conducted at 10 km x 10 km (∼0.1 degree). Although the BirdLife ranges may be different from those derived from empirical occurrence data on citizen science portals like eBird (Sullivan et al. 2009), the macroecological conclusions are known to remain robust (Aronsson et al. 2024). We also faced a challenge with the non-availability of official shapefiles (or boundaries) of many of the officially declared 1134 PAs (as of February 2025; https://wiienvis.nic.in/Database) that have only point locations available online. We sourced our PA data from the Coalition for Wildlife Corridors (CWC) website (see methods), and our assessment of the PA coverage for certain species may be limited. We also note that despite the usage of the most updated global phylogenetic tree (McTavish et al. 2025), many parts of South Asia still lack genetic data (Reddy 2014), including species found in India (132 out of 1372 species), which may limit our assessment for these species. Estimating accurate phylogenetic placements and dating estimates for species would give more robust results for evolutionary distinctiveness and other phylogeny-based metrics.

## 5. Conclusion

Our study builds on phylogenetically informed species conservation, and we reassess global avian EDGE scores and subset them to India to highlight species and areas that are threatened and evolutionarily unique. Conserving avian evolutionary history has been shown to effectively safeguard biodiversity benefits and uses for future generations (Gumbs, Gray, Hoffmann, et al. 2023).

The species with the highest EDGE score in India was the Bengal Florican, which is one of the most endangered bustards with a small and declining population attributed to the conversion of grassland habitats (Packman et al. 2014). Additionally, we find that northeastern India is exceptional in its rich assemblage of several such threatened and evolutionary unique species. However, it is also worth noting that regions like the semi-arid landscapes in western India, the Gangetic plains in north India, the Western Ghats, and the Deccan Peninsula also host a higher number of threatened and evolutionary unique species than expected based on richness patterns. We also provide EDGE scores for IBAs and PAs across India’s ten biogeographic zones to help policymakers and conservationists safeguard sites in the country with high threatened evolutionary history. IBAs of evolutionary importance were Intaki National Park (North East), Pilibhit Tiger Reserve (Gangetic Plains), Eaglenest Wildlife Sanctuary (Himalaya), Kottiyoor Reserve Forest, Brahmagiri Wildlife Sanctuary, Aralam Wildlife Sanctuary (Western Ghats), Thane Creek (Coasts), Bhimashankar Wildlife Santuary (Deccan Peninsula), Sur Sarovar Bird Sanctuary (Semi-Arid), Kangchendzonga National Park and Biosphere Reserve (Trans-Himalaya), Banni Grassland and Chhari Dhand (Desert), Mount Harriet National Park, Mahatma Gandhi Marine National Park, Kadakachang and Barangtang-Rafters Creek (Islands). Additional systematic work and updates to the phylogeny in the future are likely to change some of the EDGE scores; nevertheless, our study is the first to look at fine-scale spatial patterns of the distribution of threatened evolutionary history of birds in India to help prioritize future conservation investments.

## 6. CRediT authorship contribution statement

Archita Sharma - Conceptualization (Lead), Data Curation (Lead), Formal Analysis (Lead), Funding Acquisition (Supporting), Investigation (Lead), Methodology (Equal), Project Admin (Equal), Software (Equal), Validation (Equal), Visualization (Lead), Writing Original Draft (Lead), Writing Review & Editing (Equal)

Naman Goyal - Conceptualization (Supporting), Data Curation (Supporting), Formal Analysis (Supporting), Funding Acquisition (Supporting), Investigation (Supporting), Methodology (Equal), Software (Equal), Validation (Equal), Visualization (Supporting), Writing Original Draft (Supporting), Writing Review & Editing (Equal)

Dhyey Shah - Data Curation (Supporting), Methodology (Equal), Software (Equal), Validation (Equal) VV Robin - Conceptualization (Supporting), Funding Acquisition (Lead), Investigation (Supporting), Methodology (Equal), Project Admin (Equal), Resources (Lead), Supervision (Lead), Validation (Equal), Writing Original Draft (Supporting), Writing Review & Editing (Equal)

## 8. Acknowledgements

The study was supported by Rohini Nilekani Philanthropic Funding. We acknowledge Indian Institute of Science Education and Research Tirupati, for providing us with computational and infrastructural support. We thank Dr Prachi Thatte for sharing the Protected Area shapefiles of India and providing valuable feedback on the manuscript. We also thank Ashwin Warudkar, Varun Kher, and KL Vinay for their suggestions in improving the manuscript.

## 9. Appendix A Supplementary Figures and Table

