## Appendix A for "Using phylogenetic relationships to assess conservation priorities for birds in India"

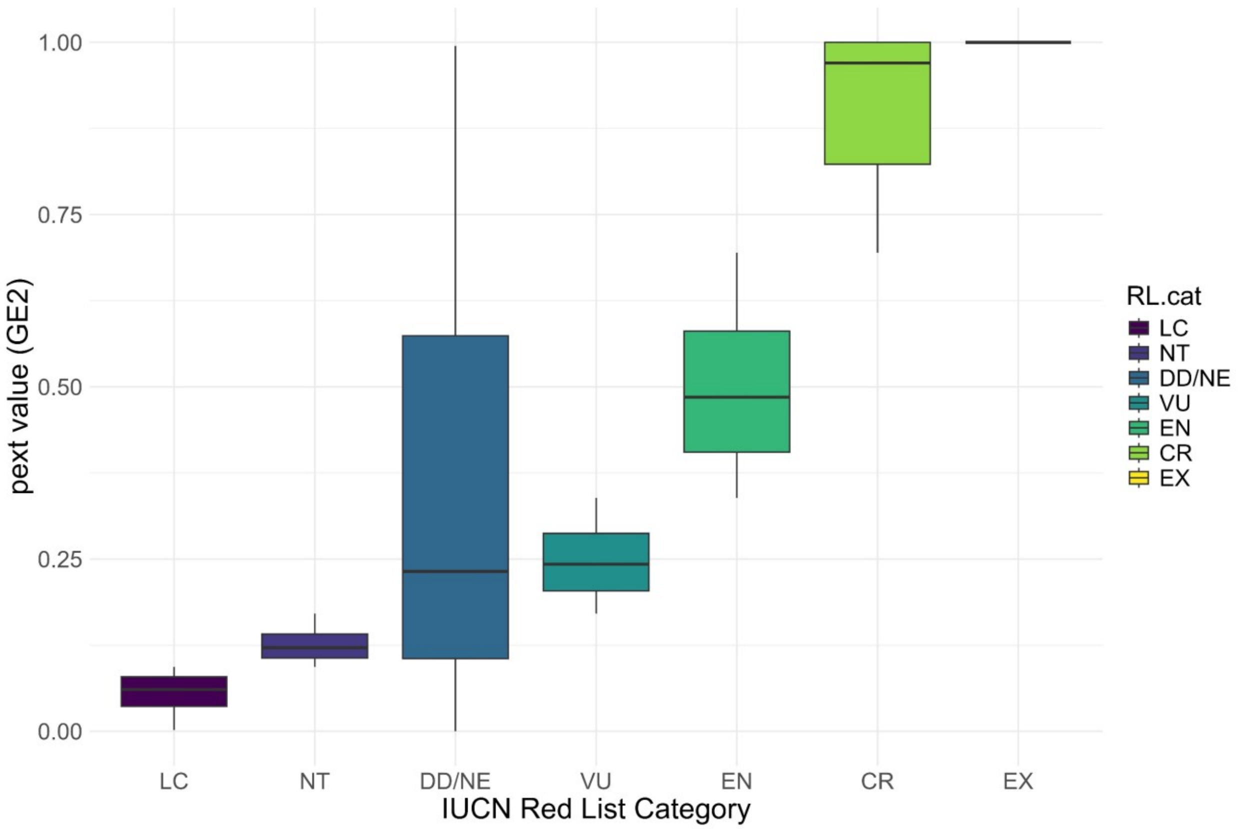


*Figure A1: Distribution of pext values (probability of extinction) for each IUCN Red List category. Functions to generate GE2 scores/pext values were downloaded from - [https://github.com/rgumbs/EDGE2/blob/main/GE.2.calc](https://github.com/rgumbs/EDGE2/blob/main/GE.2.calc). For species listed as Possibly Extinct (CR(PE)), Possibly Extinct in the Wild (CR(PEW)), and Extinct in the Wild (EW), pext is drawn from the corresponding CR pext values.*

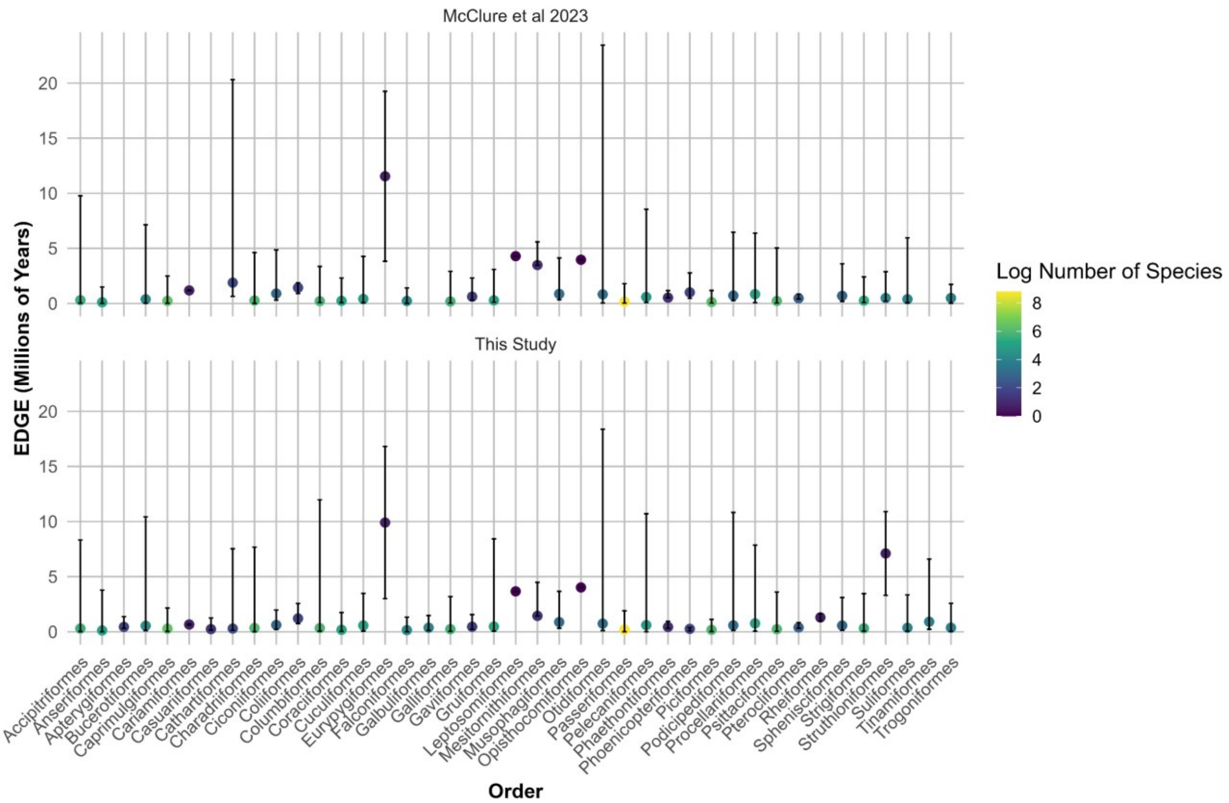


*Figure A2: Comparison of EDGE scores across global avian orders (points, median; lines, 2.5th and 97.5th percentiles) between this study and McClure et al. (2023).*

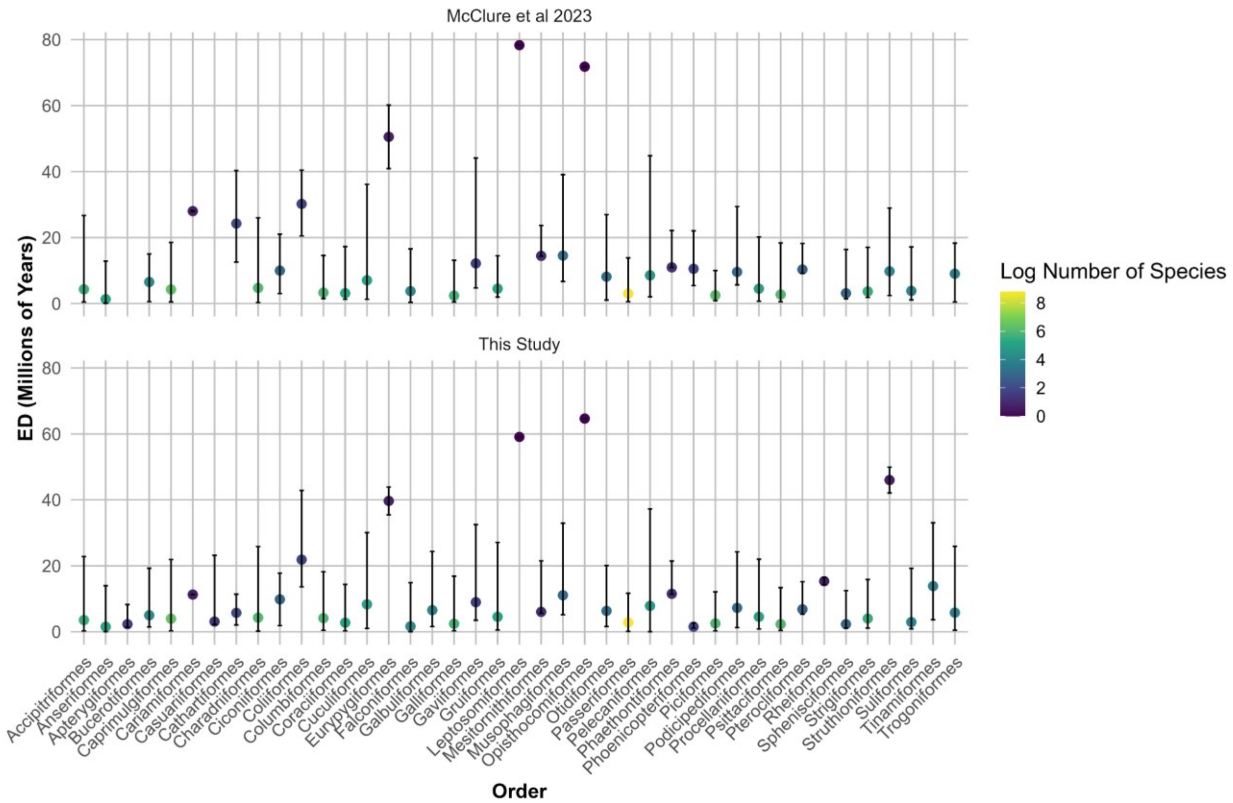


*Figure A3: Comparison of ED scores across global avian orders (points, median; lines, 2.5th and 97.5th percentiles) between this study and McClure et al. (2023).*

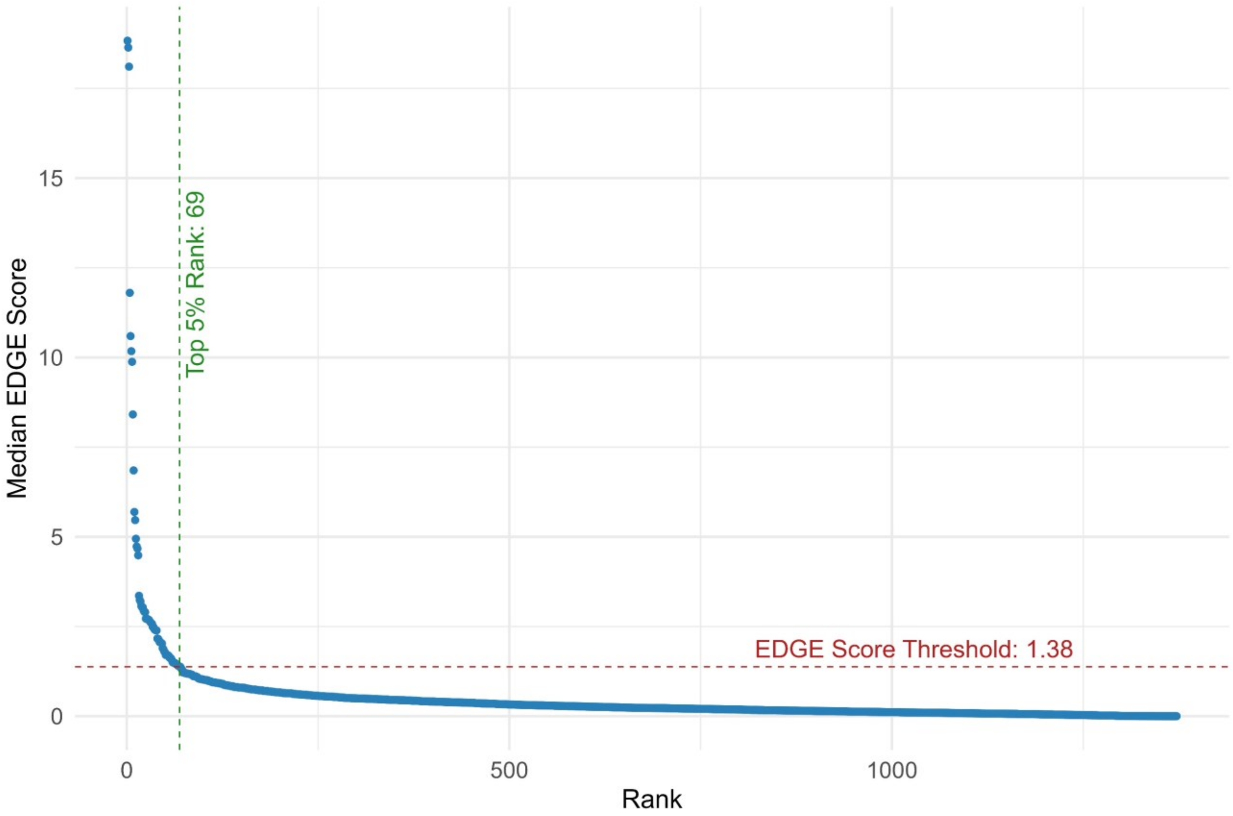


*Figure A4: EDGE score distribution and ranks across 1372 species found in India. Each point represents a species ranked in descending order of EDGE score. The dashed horizontal line indicates the 95th percentile EDGE score threshold (1.38), while the dashed vertical line denotes the rank (69) corresponding to the top 5% of species.*

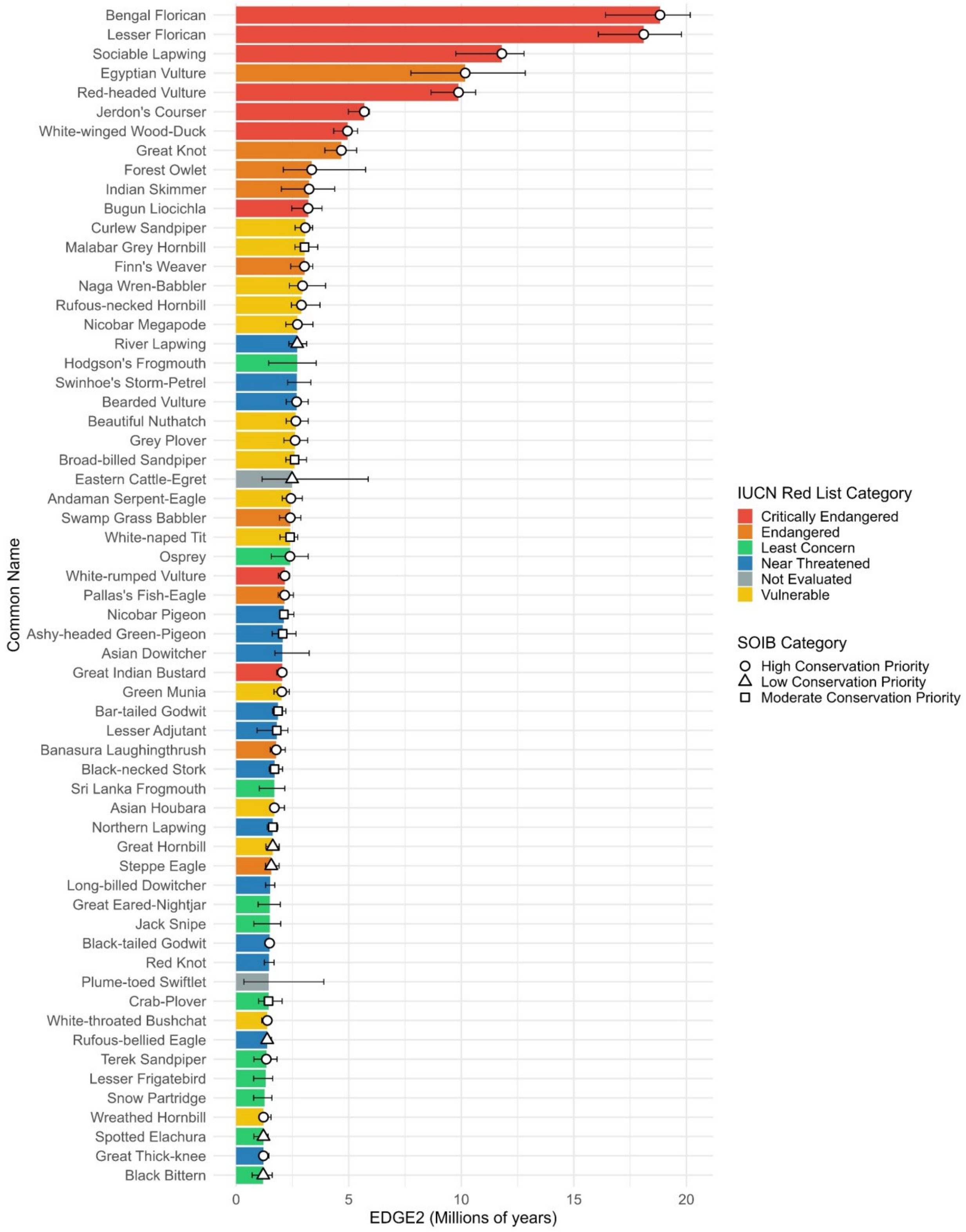


*Figure A5: Top 5% species of birds (n=61) in India with the highest median EDGE scores (removing historical and vagrant records).* Not recognized species were treated as 'Not Evaluated' for EDGE analysis.

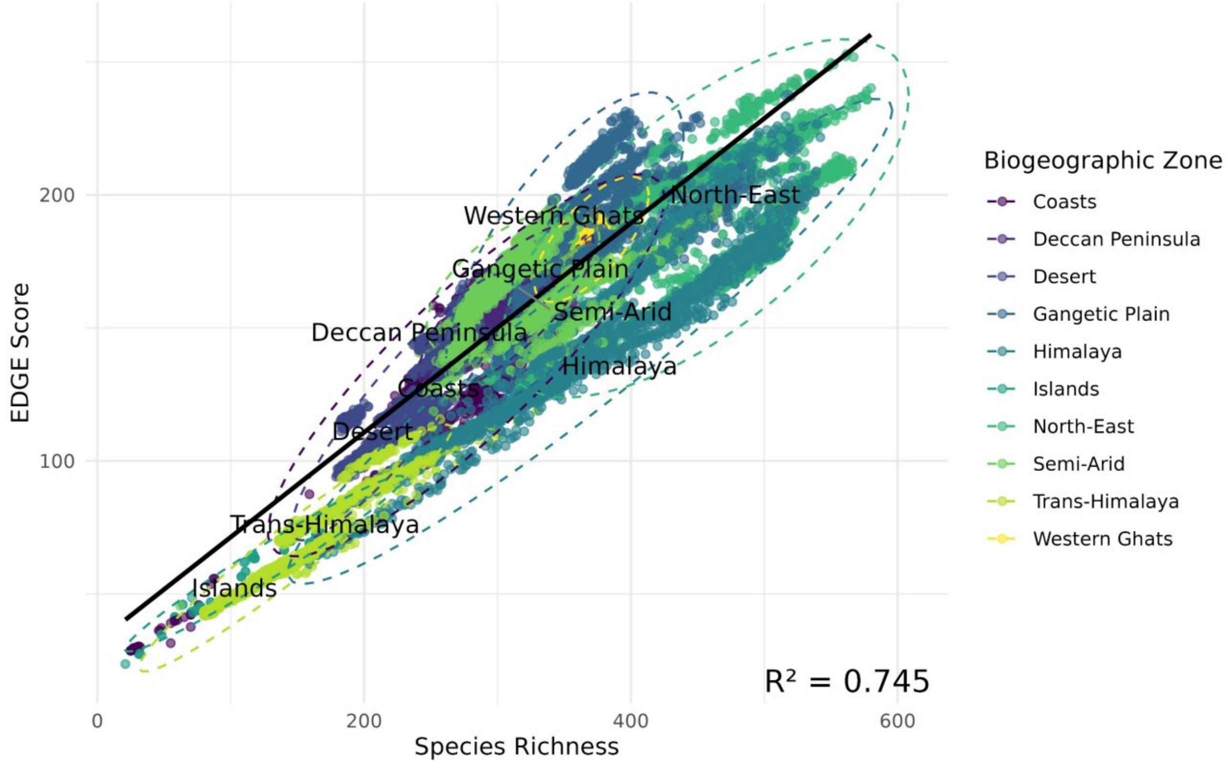


*Figure A6: Linear regression plot between EDGE score and Species Richness.*

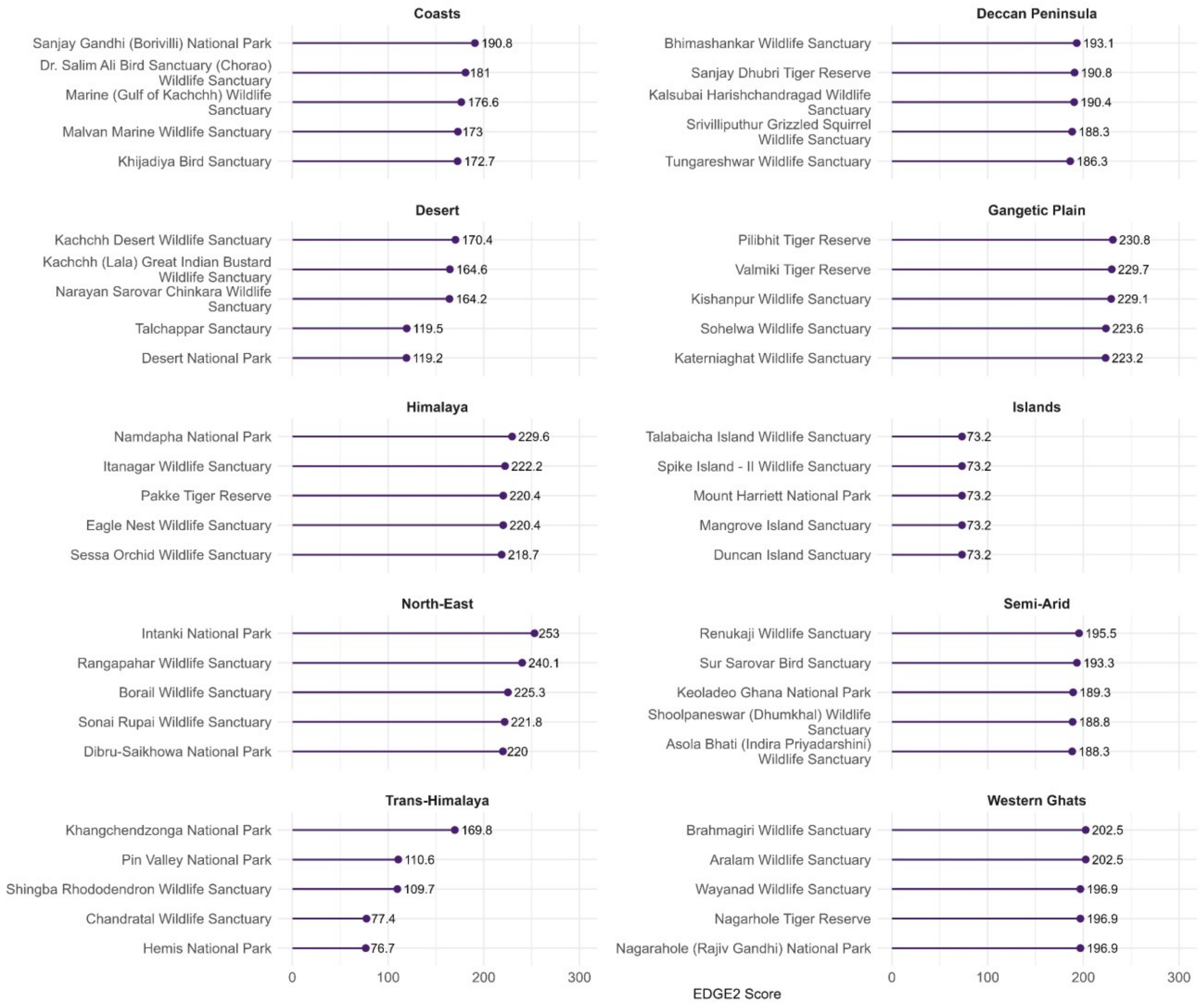


*Figure A7: Top PAs (Protected Areas) with the highest EDGE scores in each of India's 10 biogeographic zones.*

| | IUCN_Red_List_Category | Species_count_Aves | Median_pext_value |
|---|---|---|---|
| 1 | EX | 159 | 1.000000 |
| 2 | CR | 191 | 0.970000 |
| 3 | CR(PE) | 17 | 0.970000 |
| 4 | CR(PEW) | 1 | 0.970000 |
| 5 | EW | 5 | 0.970000 |
| 6 | EN | 378 | 0.485000 |
| 7 | VU | 673 | 0.242500 |
| 8 | DD | 35 | 0.232000 |
| 9 | NE | 275 | 0.232000 |
| 10 | NT | 885 | 0.121250 |
| 11 | LC | 8398 | 0.060625 |

*Table A1: IUCN Red List Categories with the respective count of species and median pext (probability of extinction) values. The taxonomy followed in this study is Clements v2023.*
